# Immunological characterization of a rat model of Duchenne’s disease and demonstration of improved muscle strength after anti-CD45RC antibody treatment

**DOI:** 10.1101/407023

**Authors:** Laure-Hélène Ouisse, Séverine Remy, Aude Lafoux, Thibaut Larcher, Laurent Tesson, Vanessa Chenouard, Carole Guillonneau, Lucas Brusselle, Nadège Vimond, Karl Rouger, Yann Péréon, Alexis Chenouard, Christèle Gras-Le Guen, Cécile Braudeau, Régis Josien, Corinne Huchet, Ignacio Anegon

## Abstract

Duchenne muscular dystrophy (DMD) has as standard pharmacological therapy with corticoisteroids (CS) that decrease inflammation and immune responses present in patients and animal models. CS have however limited efficacy and important and numerous side effects. Therefore, there is a need for new anti-inflammatory and pro-tolerogenic treatments that could replace or decrease doses of CS. We first assessed the status of immune system of dystrophin-deficient rats (*Dmd*^*mdx*^) that closely reproduce the phenotype of DMD patients. *Dmd*^*mdx*^ rats showed increased leukocyte infiltration in skeletal and cardiac muscles, containing mostly macrophages but also T cells, and increased expression of several cytokines. Anti-CD45RC Monoclonal antibody (Mab) treatment induced immune tolerance in models of organ transplantation and GVHD (Graft Versus Host Disease). We observed that muscles and blood of DMD patients contained T CD4^+^ and CD8^+^ expressing high levels of CD45RC^high^ cells. Treatment of young *Dmd*^*mdx*^ rats with anti-CD45RC MAb corrected skeletal muscle strength associated to a depletion of effectors CD45RC^high^ T cells with no obvious side-effects. Prednisolone treatment of *Dmd*^*mdx*^ rats similarly increased skeletal muscle strength and was also associated to a depletion of effectors CD45RC^high^ cells but resulted in severe weight loss.

Overall, *Dmd*^*mdx*^ rats display important immune inflammatory response and thus represent a useful model to analyze new anti-inflammatory and tolerogenic treatments for DMD. As an example, a new treatment with anti-CD45RC antibodies improved muscle strength in *Dmd*^*mdx*^ rats as prednisolone did but without side effects. Anti-CD45RC therapy could complement other therapies in DMD patients.

## Introduction

Duchenne muscular dystrophy (DMD) is the most common inherited muscle disease. It is caused by a mutation in the dystrophin gene with a X-chromosomal recessive inheritance that affects 1 of 3,500 male births ^1^. It has a severe prognosis with life expectancy ranging from the late teens to the mid-30s. Muscle fibers show necrosis and regeneration/degeneration associated to inflammation with progressive replacement by connective and adipose tissue ^1^. The mdx mouse carries a mutation in the *Dmd* gene and is a well-established mouse model of DMD. Nevertheless, the muscle impairment is rather mild in *mdx* mice compared with DMD patients indicating that new animal models are required ^2^.

We previously generated *Dmd-*deficient (*Dmd*^*mdx*^) rats using TALENs ^3^. Forelimb and hindlimb muscular strength and spontaneous activity were decreased. Skeletal and cardiac muscles showed necrosis and regeneration of muscle fibers associated to progressive replacement by fibrotic and adipose tissue. Weak muscle strength and muscle lesions therefor closely mimic those observed in DMD patients. *Dmd*^*mdx*^ rats represent a useful small animal model of pre-clinical research for DMD ^4^.

To date, there is no cure for muscular dystrophies, and despite that gene and cell therapies will likely bring in the future cure of the disease there is still need for therapies for associated pathology such as immune responses and inflammation. Immune responses are involved in the pathophysiology of disease in both DMD patients and *mdx* mice [for a review see ^5^]. Standard therapy of DMD is based on treatment with corticosteroids (CS), which have been shown to act at least in part through anti-inflammatory actions and inhibition of CD8^+^ T cells that improve muscle strength in a fraction of patients ^5-7^. Apart from its moderate efficacy, CS treatment is limited by serious systemic side effects, such as short stature, obesity, psychological symptoms, osteoporosis, diabetes and hypertension ^6^. Furthermore, CS through their broad and nonspecific anti-inflammatory effects inhibit inflammatory mechanisms that promote muscle repair ^5^.

T effector cells against DMD have been described in patients before and after gene therapy ^8-10^ CD4^+^ T regulatory cells (Tregs) limit the severity of the disease in *mdx* mice not only through inhibition of immune responses but also by their tissue repair activity ^5, 11, 12^.

Thus, inhibition of immune responses and promotion of immune tolerance are potentially important adjuvants to the therapeutic arsenal to treat DMD patients but these immunointerventions should at the same time preserve immune responses that promote muscle regeneration as well as protection against pathogens and cancer cells. Knowledge of immune responses in DMD patients and animal models are thus important for targeted immunointerventions associated to other treatments such as gene or cell therapy. Furthermore, immune responses can also be an obstacle to gene and cell therapy since in both situations newly produced dystrophin could be recognized as immunogenic and cells expressing it destroyed ^10^. Thus, analyses of immune cells and immunotherapies in *Dmd*^*mdx*^ rats could give potentially important results for development of new treatments for DMD patients.

We have described that CD4^+^ and CD8^+^ Tregs in rats and humans are comprised within CD45RC^low/-^ cells ^13, 14^. We have recently shown that treatment with anti-CD45RC monoclonal antibody (MAb) in a rat model of allograft rejection and in mouse immune humanized models of graft versus host disease (GVHD) could induce permanent allograft acceptance and inhibition of GVHD ^14^. Anti-CD45RC treatment depleted only T cells that were CD45RC^high^, i.e. naïve T cells, precursors of Th1 cells and effector memory T cells including TEMRA cells, whereas CD8^+^ or CD4^+^ Tregs, both in rats and humans, are CD45RC^low/-^ and thus were spared. Among CD45RC^low/-^ cells, CD8^+^ and CD4^+^ Tregs specific for donor alloantigens protect from graft rejection. Importantly, immune responses against third party donors and exogenous antigens were preserved, thus anti-CD45RC antibody treatment does not result in broad immunosuppression but rather specific elimination of T cells with effector functions and preserved Tregs followed by their activation and expansion 14.

We thus reasoned that treatment of *Dmd*^*mdx*^ rats with anti-CD45RC MAbs could eliminate CD45RC^high^ effector T cells and enrich CD45RC^low/-^ Tregs. The later could then act at the same time as inhibitors of immune responses and favoring muscle repair and homeostasis. To the best of our knowledge, treatment with antibodies directed against other cell antigens that favor immune tolerance in transplantation, GVHD or autoimmune diseases, such as anti-CD3, -CD28, -CD127 or -CD137, have not been reported in none of the other animal models of DMD.

We first analyzed immune parameters in *Dmd*^*mdx*^ rats and we secondly treated *Dmd*^*mdx*^ rats with the same anti-CD45RC MAb previously used to induce allograft tolerance in comparison to the standard of care (i.e. prednisolone). We observed that the skeletal and cardiac muscle of *Dmd*^*mdx*^ rats showed a leukocyte infiltrate predominantly formed by macrophages and to a lesser extent by T cells. M2 type macrophages increased with time. Treatment with an anti-CD45RC depleting MAb resulted in increased muscle strength associated to a decrease in T cells but not of macrophages. Prednisolone treatment also increased muscle strength and decreased CD45RC^high^ cells but decreased growth of *Dmd*^*mdx*^ rats whereas anti-CD45RC did not. CD45RC^+^ cells are also present in the blood and muscles of DMD patients.

Overall, immune responses and inflammation are present in the *Dmd*^*mdx*^ rat muscles and anti-CD45RC MAb treatment resulted in amelioration of skeletal muscle strength. This is the first report showing that a treatment with a monoclonal antibody targeting specific T cell populations results in amelioration of clinical parameters in a pre-clinical model of DMD.

## Results

### Increased mononuclear leukocyte infiltration in skeletal muscles of *Dmd*^*mdx*^ rats

Mononuclear leukocytes in the muscle and spleen of *Dmd*^*mdx*^ rats were analyzed by flow cytometry using the pan-leukocyte marker CD45 **(Fig 1).** The number of total mononuclear leukocytes in the muscle of littermate wild type (WT) and *Dmd*^*mdx*^ rats were comparable at 2 weeks of age, but at 4 weeks *Dmd*^*mdx*^ rats showed a sharp increase that was maintained until week 8 and then decreased at weeks 12 and 14 to values that were still significantly higher than those observed in littermate WT rats **(Fig. 1A-B)**. Granulocytes were rarely observed at early time points in biopsies stained with Hemalun-Eosin-Saffron (data not shown). Total leukocyte numbers in the spleen were comparable between WT and *Dmd*^*mdx*^ rats at all-time points analyzed **(Fig. 1A)**. Thus, limb muscles of *Dmd*^*mdx*^ rats showed an anatomical specific leukocyte infiltrate that indicates the presence of a localized immune/inflammatory response.

**Figure 1.**
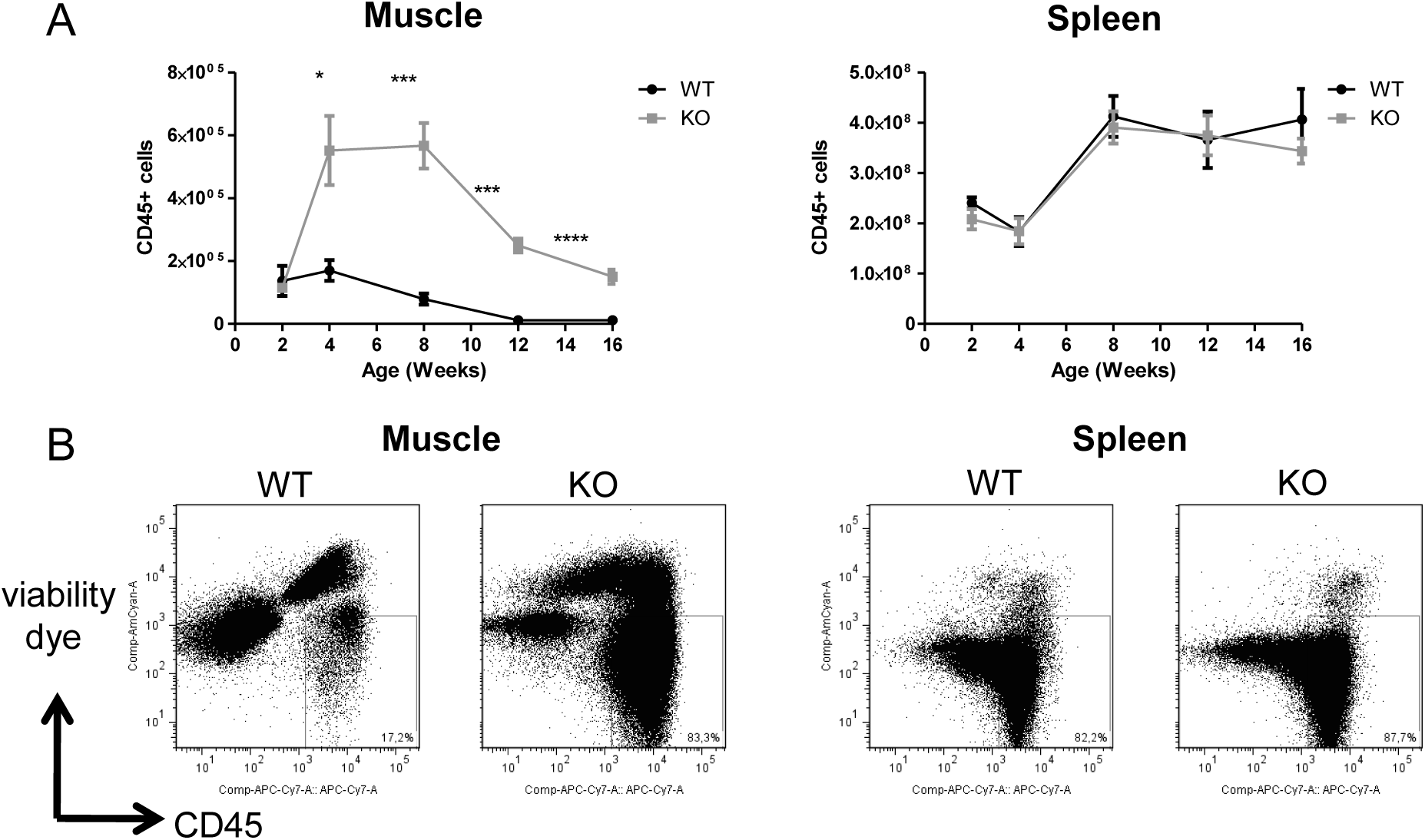
Number of leukocytes in skeletal muscle and spleen of *Dmd*^*mdx*^ rats. Hind limb muscles and spleen were harvested from littermate wild-type (WT) or *Dmd*^*mdx*^ (KO) rats at the indicated time points of age. Muscles and spleens were digested with collagenase, mononuclear cells were isolated using a density gradient and analyzed by cytofluorimetry. **A)** Number of viable CD45^+^ cells per gram of muscle (left panel) or whole spleen (right panel) at different time points. WT, n=4, 5, 7, 7, 9 at 2, 4, 8, 12 and 16 weeks, respectively; *Dmd*^*mdx*^, n=3, 6, 10, 11, 16, at 2, 4, 8, 12 and 16 weeks, respectively * p< 0.05, ** p< 0.01, and *** p< 0.001. **B)** Representative dot-plot analysis of viable SSC CD45^+^ mononuclear leukocytes from muscle (left panel) or spleen (right panel) from animals at 12 weeks of age.

### Presence of macrophages and T cells in skeletal muscle of *Dmd*^*mdx*^ rats as analyzed by cytofluorimetry

We used flow cytometry analysis to obtain frequencies and absolute numbers, of different mononuclear leukocyte populations. The analysis of viable CD45^+^ mononuclear leukocyte subpopulations showed that ~90% of muscle infiltrating cells in *Dmd*^*mdx*^ rats were CD68^+^ (vs. ~60% in WT rats), increasing sharply at 4 weeks, maximal at 8 weeks and decreased but were still higher than WT at 12 and 16 weeks of age and were of higher granularity as shown by their SSC profile **(Fig. 2A).** In contrast, numbers of spleen CD68^+^ macrophages increased steadily with age and were comparable between *Dmd*^*mdx*^ and WT rats **(Fig. 2A).** Identical results were obtained with the macrophage marker SIRPα **(Supplementary figure 1).**

**Figure 2.**
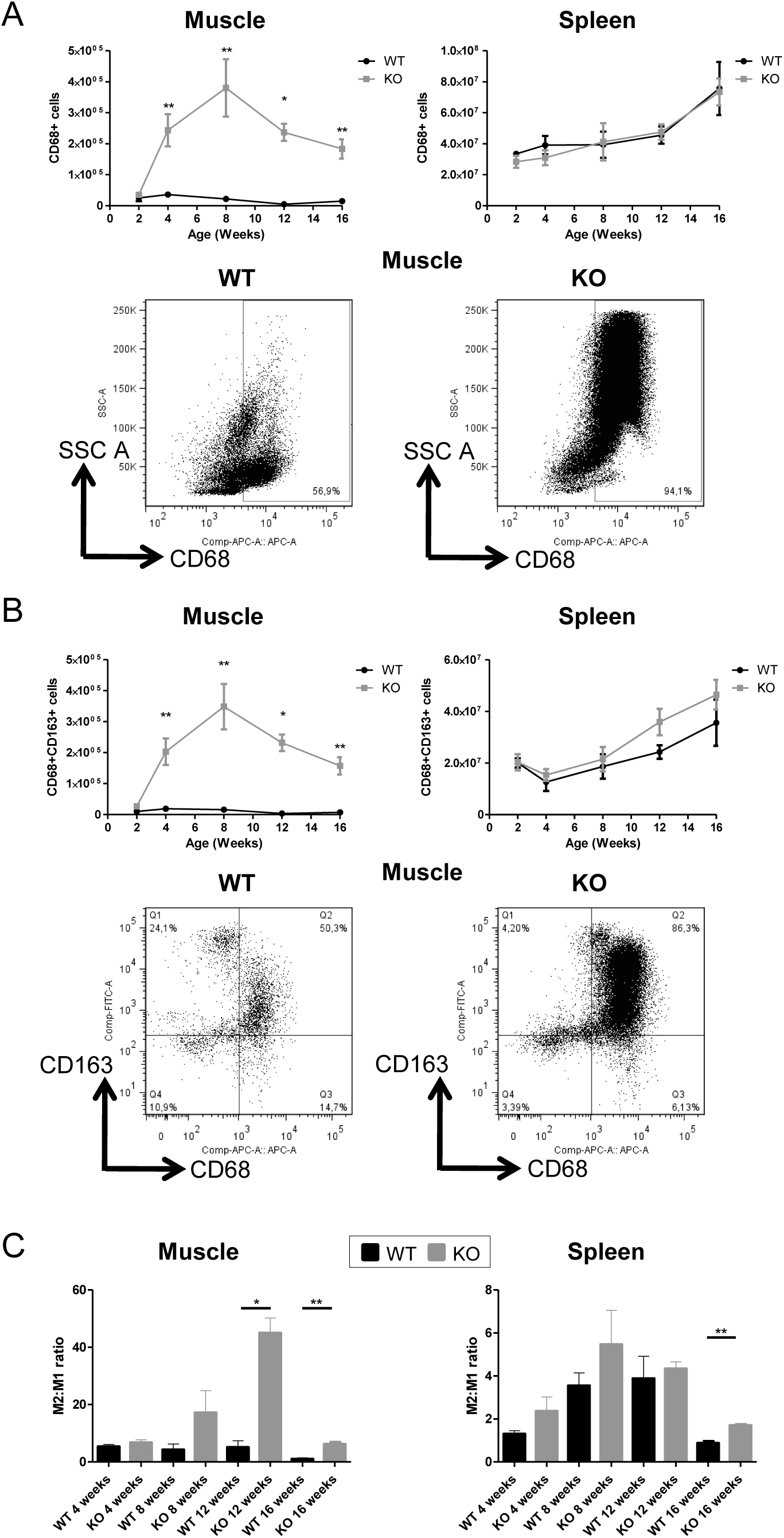
Macrophages in skeletal muscle and spleen of *Dmd*^*mdx*^ rats. Cytofluorimetry of single-cell suspensions from hind limb muscles or spleen WT or *Dmd*^*mdx*^ (KO) at the indicated time points of age. **A)** Total number of macrophages CD68^+^ cells per gram of muscle (upper left panel) or of whole spleen (upper right panel). Representative dot plots of macrophages high granularity using side scatter (SSC^high^) CD68^+^ cells after gating on viable (negatively-stained cells) CD45^+^ cells from muscle of WT or *Dmd*^*mdx*^ 12 weeks-old rat**s** (lower panel). **B)** Total number of viable CD68^+^CD163^+^ type 2 macrophages per gram of muscle (upper left panel) or whole spleen (upper right panel). Representative dot plots of viable CD68^+^CD163^+^ cells from muscle of WT or *Dmd*^*mdx*^ 12 weeks-old rat**s** (lower panels). **C)** Macrophages type 2 (CD68^+^CD163^+^) over type 1 macrophages (CD68^+^CD163^-^) ratios in muscle (left panel) or spleen (right panel) of WT (black) or *Dmd*^*mdx*^ (grey) rats. n=3, 6, 6, 7 and 8 (at 2,4, 8, 12 and 16 weeks of age, respectively) for *Dmd*^*mdx*^ rats and n= 4, 6, 4, 3 and 4 (at 2,4, 8, 12 and 16 weeks of age, respectively) for WT rats. * p< 0.05, *** p< 0.01. Results were obtained from several experiments performed using all groups of animals in each experiment.

Analysis of the M2 marker CD163 also showed a similar curve with an increase at 4 weeks, maintained at 8 weeks and a decrease at 12 and 16 weeks of age with an increase in CD68 expression levels in some animals **(Fig. 2B)**. In contrast, the number of CD163^+^ macrophages in spleen increased with age and were comparable between *Dmd*^*mdx*^ rats and WT rats **(Fig. 2B)**. The ratio of M2:M1 macrophages in muscles of *Dmd*^*mdx*^ rats was comparable at 4 weeks, increased non significantly at 8 weeks and was significantly higher at 12 and 16 weeks of age, whereas it was constant in muscles of WT rats **(Fig. 2C).** In the spleen, ratio of M2:M1 macrophages increased with time but was always lower than in muscles and comparable for both *Dmd*^*mdx*^ and WT rats except at 16 weeks of age in which *Dmd*^*mdx*^ showed a modest but significant increase vs. WT rats **(Fig. 2C).**

Analysis of T cells in muscles showed that total TCR^+^αβ cells **(Fig. 3A-B)** as well as CD4^+^ **(Fig.3 C-D)** and CD8^+^ T cells **(Fig. 3E-F)** in *Dmd*^*mdx*^ rats increased sharply at 4 weeks and then decreased at later time points with significantly higher levels at 4 and 12 weeks vs. WT animals. In contrast, in the spleen, total TCR^+^ cells, CD4^+^ T cells and CD8^+^ T cells for both *Dmd*^*mdx*^ and WT rats increased steadily to comparable numbers at 8 weeks and remained stable **(Fig. 3A-F)**.

**Figure 3.**
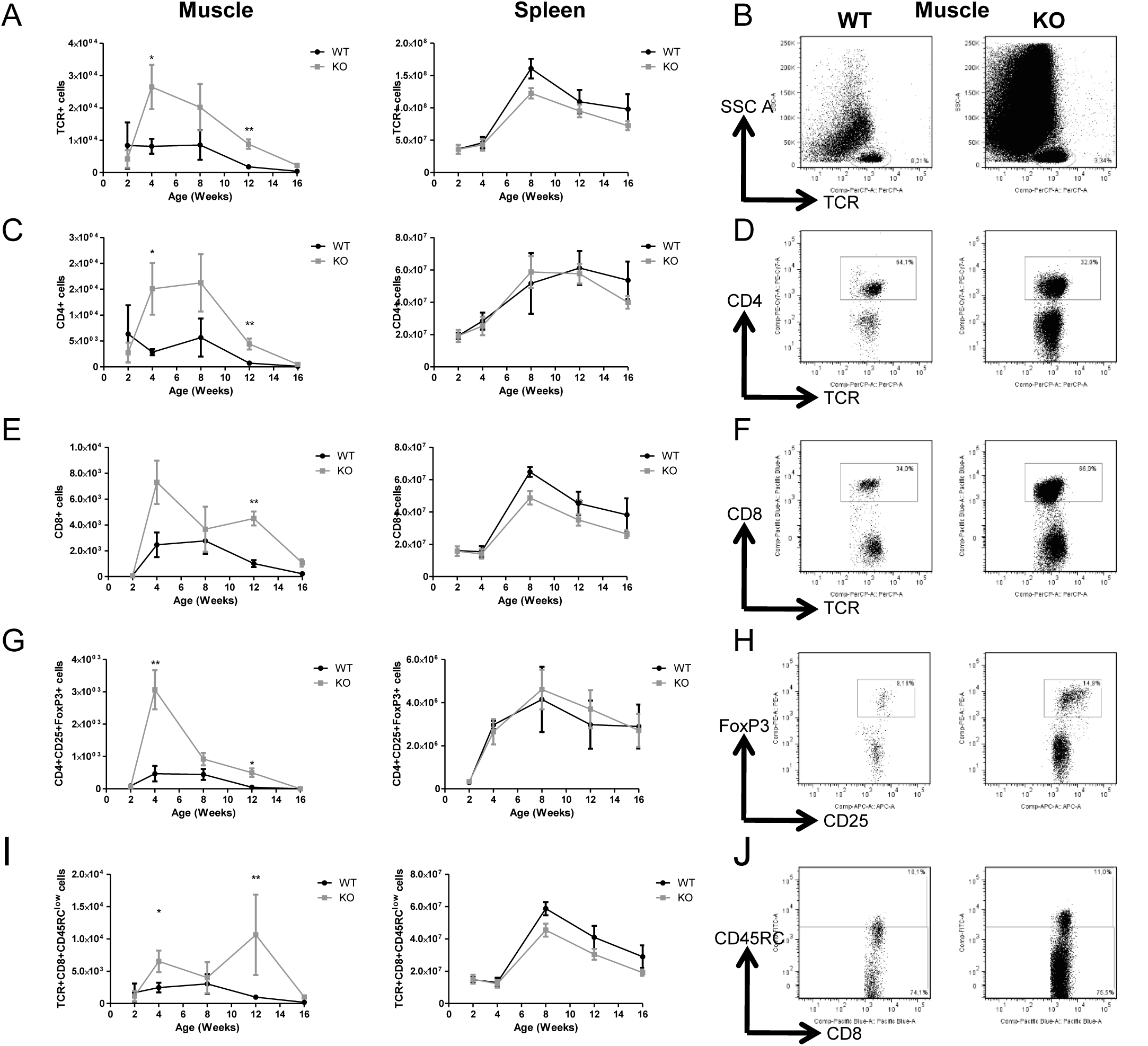
T cells in skeletal muscle and spleen of *Dmd*^*mdx*^ rats. Hind limb muscles or spleen from WT or *Dmd*^*mdx*^ (KO) at the indicated time points of age were harvested, collagenase digested and analyzed by cytofluorimetry. **A)** Total numbers of viable CD45^+^TCR^+^ cells per gram of muscle (left panel) and of total spleen (right panel). **B)** Representative dot plots of viable CD45^+^TCR^+^ cells from muscle of WT or *Dmd*^*mdx*^ 12 weeks-old rats. **C)** Total number of CD45^+^TCR^+^CD4^+^ cells per gram of muscle (left panel) and of total spleen (right panel). **D)** Representative dot plots of WT or *Dmd*^*mdx*^ 12 weeks-old rat muscle single-cell suspension showing gating on viable CD45^+^TCR^+^CD4^+^ cells. **E)** Total number of TCR^+^CD8^+^ cells per gram of muscle (left panel) and of total spleen (right panel). **F)** Representative dot plots of WT or *Dmd*^*mdx*^ 12 weeks-old rat muscle single-cell suspension showing gating on viable CD45^+^TCR^+^CD8^+^ cells. **G)** Total number of TCR^+^CD4^+^CD25^+^Foxp3^+^ cells per gram of muscle (left panel) and whole spleen (right panel). **H)** Representative dot plots of WT or *Dmd*^*mdx*^ 12 weeks-old rat muscle single-cell suspension showing gating on viable CD45^+^TCR^+^CD4^+^CD25^+^Foxp3^+^ cells. **I)** Total number of TCR^+^CD8^+^CD45RC^low/--^ cells per gram of muscle (left panel) and whole spleen (right panel). **J)** Representative dot plots of WT or *Dmd*^*mdx*^ 12 weeks-old rat muscle single-cell suspension showing gating on viable CD45^+^TCR^+^CD8^+^CD45RC^low/-^ cells. n=3, 6, 10, 12 and 4 (at 2, 4, 8, 12 and 16 weeks of age, respectively) for *Dmd*^*mdx*^ rats and n= 4, 5, 7, 7 and 4 (at 2, 4, 8, 12 and 16 weeks of age respectively) for WT rats. Results were obtained from several experiments performed using all groups of animals in each experiment.

The muscles of *Dmd*^*mdx*^ vs. WT rats showed significantly increased levels of Foxp3^+^ CD4+ Tregs at 4 and 12 weeks **(Fig. 3G-H)**. As we previously described ^15, 16^, CD8^+^ Tregs were defined as CD8^+^CD45RC^low^ T cells and were significantly increased at 4 and 12 weeks **(Fig. 3I-J)**. In contrast, in spleen, total Foxp3^+^CD4+ Tregs and CD8^+^CD45RC^low^ Tregs of both *Dmd*^*mdx*^ and WT rats increased comparably at 8 weeks and remained stable **(Fig. 3H and J)**.

B cells (CD45RA^+^ and CD45R^+^) and NK cells (CD161^high^) represented respectively always <2% and 3% of total muscle leukocytes in both *Dmd*^*mdx*^ and WT rats and were comparable in spleens of both *Dmd*^*mdx*^ and WT rats **(data not shown).**

Thus, the majority of leukocytes in muscles of *Dmd*^*mdx*^ rats were macrophages that reached maximal levels between 4 and 8 weeks of age and the ratio of M2:M1 increased at 12 and 16 weeks of age. T cells, including CD8^+^ and CD4^+^ Tregs, showed a similar evolution with similar proportions of both CD4^+^ and CD8^+^ cells.

### Detection of macrophages in cardiac and skeletal muscle of *Dmd*^*mdx*^ rats as analyzed by immunohistology

We used tissue immunofluorescence to analyze leukocyte populations in cardiac muscle since flow cytometry analysis required numbers of cells higher than we could routinely obtain from hearts from WT origin and to confirm the presence in skeletal muscle of leukocytes defined by flow cytometry. Skeletal and cardiac muscle biopsies at 8 and 12 weeks of age showed the presence of CD68^+^ and CD163^+^ macrophages but few CD3^+^ cells in connective tissue of both skeletal and cardiac muscles of *Dmd*^*mdx*^ rats. In comparison, only a few CD68+ macrophages were observed sporadically in WT rats **(Fig. 4).** CD163^+^ macrophages were notably numerous in foci of mononuclear cell infiltration in the cardiac muscle. Thus, immunohistology of skeletal muscles from *Dmd*^*mdx*^ rats confirmed results obtained by flow cytometry and revealed very similar pattern in cardiac muscle. As previously described ^3^, increased fibrosis **(Fig. 4)**, fiber necrosis and regeneration **(data not shown)** are present in skeletal and cardiac muscle of *Dmd*^*mdx*^ as soon as 4 weeks and more severely at 8 weeks of age. Along with these lesions, total creatinine kinase (CK) levels in serum, released from damaged muscle fibers, were comparable at week 2 between *Dmd*^*mdx*^ and WT rats and then increased significantly in *Dmd*^*mdx*^ rats to reach peak levels between weeks 4 and 8 and decreased at 12 weeks, returning to normal levels at week 16 **(Supplementary figure 2).**

**Figure 4.**
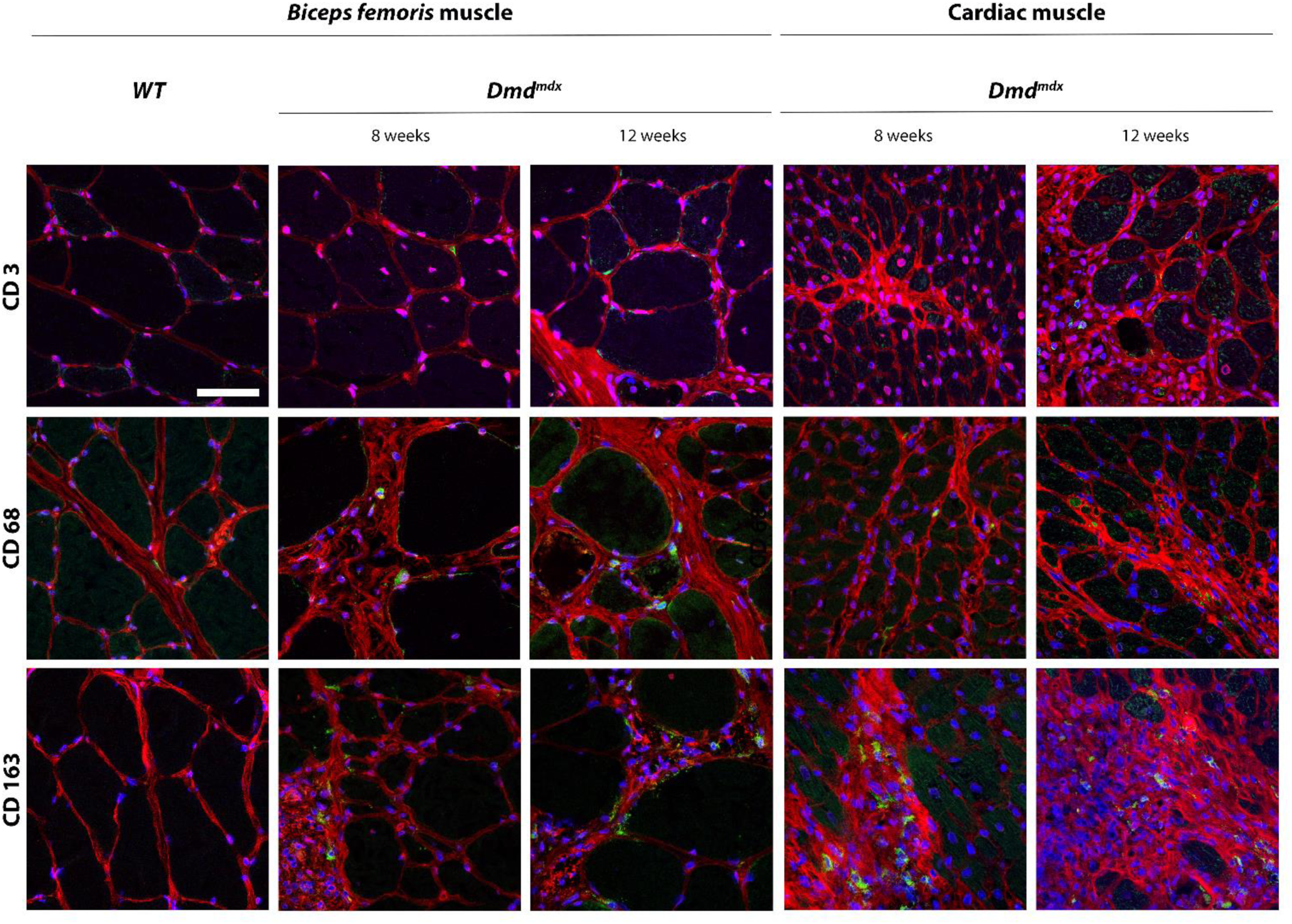
Immunohistochemical detection of leukocytes in skeletal and cardiac muscle of *Dmd*^*mdx*^ rats. Skeletal muscle (*Biceps femoris*) and cardiac muscle were harvested at 8 and 12 weeks of age from wild-type (WT) and *Dmd*^*mdx*^ (KO) rats. **A)** Tissue sections were stained with Draq5 to label nuclei (blue), with wheat germ agglutinin for connective tissue (red) and with MAbs for detection of cells expressing CD3, CD68 or CD163 (green). Scale bar identical for all pictures: 100 µm.

These results indicate that infiltration of muscle by leukocytes was associated to damaged muscle fibers and elevated CK serum levels.

### Inflammatory and growth factors in leukocytes infiltrating muscle and serum of *Dmd*^*mdx*^ rats

We used quantitative RT-PCR to analyze mRNA levels of several molecules involved in the initiation or suppression of immune responses and inflammation, as well as some muscle trophic factors in isolated mononuclear leukocytes from muscles of *Dmd*^*mdx*^ and WT at 8 and 12 weeks of age **(Fig. 5A).** TNFα expression was particularly and strongth increased in mononuclear cells from muscles of *Dmd*^*mdx*^ vs. WT at 8 weeks. Similarly, heme oxygenase-1 (HO-1), IFNγ, TGFβ, IL-10 as well as the muscle trophic factor amphiregulin ^11^ were significantly increased at 8 and/or 12 weeks **(Fig. 5A).** Arginase and IL-34 were decreased in mononuclear cells from muscle of *Dmd*^*mdx*^ rats vs. WT rats, at weeks 8 and 12 respectively **(Fig. 5A)**. IL-6 and iNOs were not statistically different in *Dmd*^*mdx*^ vs. WT rats but the former showed higher numerical levels in *Dmd*^*mdx*^ rats **(Fig. 5A).** Relaxin3 and indoleamine 2,3-dioxygenase (IDO) were detectable at very low levels without differences among the different groups of animals **(data not shown).**

**Figure 5.**
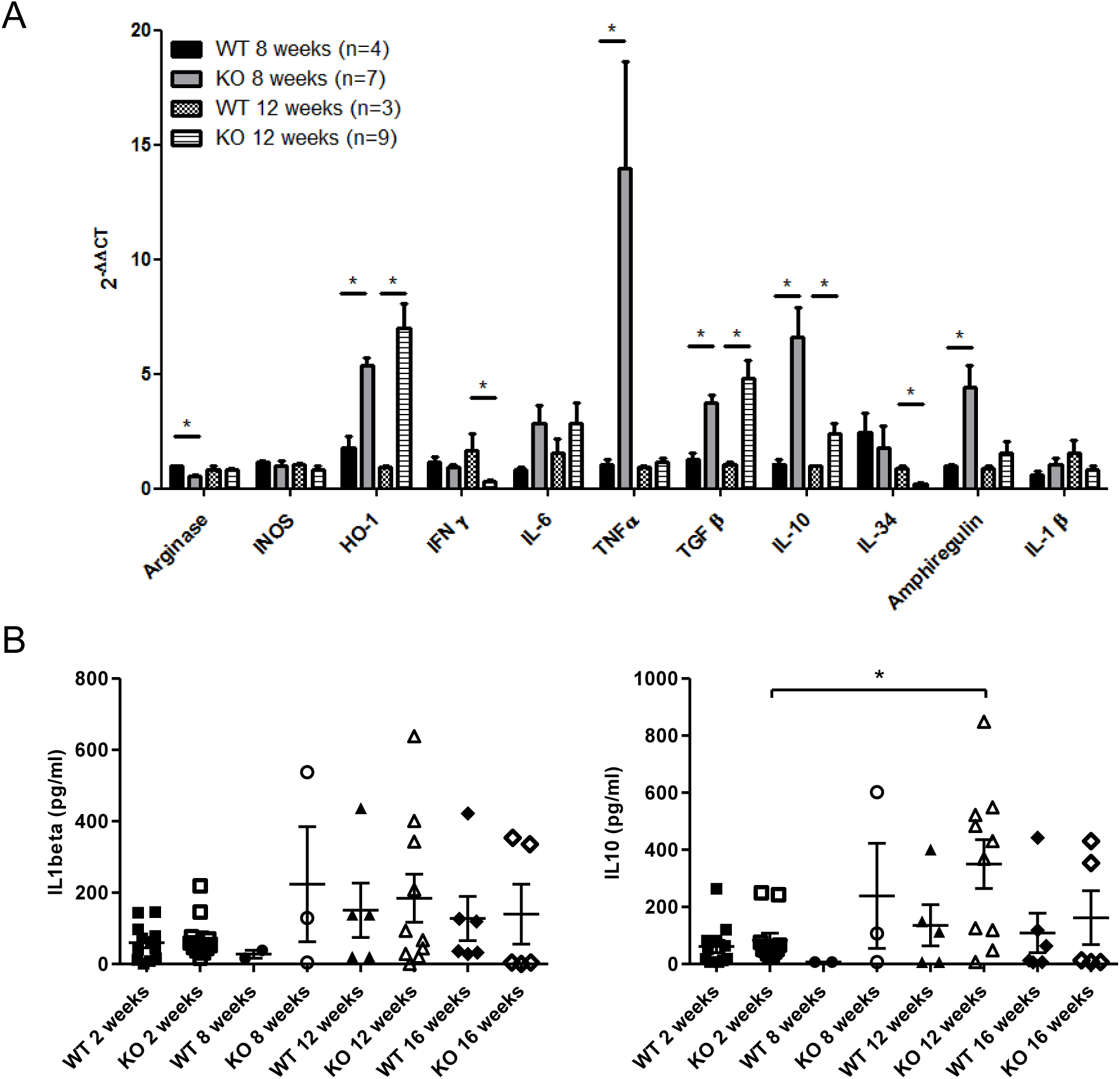
Inflammation markers and growth factors in skeletal muscle of *Dmd*^*mdx*^ rats. **A)** Mononuclear cells from skeletal muscles were harvested at 8 and 12 weeks of age from wild-type (WT) and *Dmd*^*mdx*^ (KO) rats. Total RNA was extracted and mRNA levels for the indicated molecules were analyzed by quantitative RT-PCR. * p<0.05. **B)** IL1β (left panel) and IL10 (right panel) levels in the sera of *Dmd*^*mdx*^ (n=11, 3, 10, 5 at 2, 8, 12 and 16 weeks of age, respectively) or WT (n= 12, 2, 5, 6 at 2, 8, 12 and 16 weeks of age, respectively) rats. * p<0.05.

To further evaluate cytokines in *Dmd*^*mdx*^ rats, we analyzed by using a multiplex assay the presence of cytokines in the sera of animals at different time points. IL-1beta and IL-10 were detectable in serum in both *Dmd*^*mdx*^ and WT rats without significant differences between them at 8, 12 and 16 weeks and as compared to 2 weeks IL-10 was significantly elevated only at 12 weeks **(Fig. 5B).** TNFα and IL-6 levels were undetectable **(data not shown).**

Overall, several mediators of inflammation were increased in muscle or serum, such as TNFα and IL-1β, respectively, and several anti-inflammatory molecules, such as HO-1, TGFβ, amphiregulin and IL-10 were increased in muscle, as well as the later also in serum.

### Treatment with anti-CD45RC MAb depleted CD45RC^high^ T cells and improved skeletal muscle strength

Anti-CD45RC MAb treatment induces organ transplantation tolerance and inhibits GVHD at least partially mediated by depletion of T CD8^+^CD45RC^high^ and CD4^+^CD45RC^high^ cells involved in organ rejection and GVHD and in the organ transplantation model by increased suppressor activity against donor antigens by CD8^+^CD45RC^low/-^ and CD4^+^CD45RC^low/-^ Tregs ^14^. Since CD45RC expression levels can differ in different rat strains ^17^ and have not been reported in muscle, we first analyzed the distribution of CD45RC^high^ and CD45RC^low/-^ leukocytes within different leukocytes subsets in the muscle and spleen of *Dmd*^*mdx*^ and WT Sprague-Dawley rats.

In muscles of *Dmd*^*mdx*^ rats, we observed that, within the CD8^+^ T cell population, absolute numbers of CD45RC^low/-^ **(Fig. 3I-J)** and CD45RC^high^ cells **(Supplementary figure 3A)** increased sharply and significantly in *Dmd*^*mdx*^ vs. WT at 4 weeks, remained elevated at 8 weeks and then decreased at 12 weeks to low levels observed in WT rats. Numbers in spleen of CD45RC^high^ and CD45RC^low/-^ cells were comparable of both *Dmd*^*mdx*^ and WT rats **(Supplementary figure 3A and Fig. 3I).**

For the TCR^+^CD4^+^ cell population absolute numbers of CD45RC^low/-^ cells increased significantly at 4 and 12 weeks in the muscle of *Dmd*^*mdx*^ rats vs. WT and in the spleen increased progressively without statistical differences **(Supplementary figure 3C-D).**

Absolute numbers of CD45RC^high^ cells in muscle of *Dmd*^*mdx*^ rats increased but not significantly vs. WT rats and in the spleen there were no differences between *Dmd*^*mdx*^ and WT rats **(Supplementary figure 3E-D).**

For the non-T cells, which were mostly macrophages, CD45RC^low/-^ increased significantly at 4 weeks, remained elevated at 8 weeks and decreased at 12 weeks. **(Supplementary figure 3F-G).** TCR^-^ CD45RC^high^ increased non significantly at 4 and 8 weeks, decreased at weeks 12 and 16 and are statistically higher in *Dmd*^*mdx*^ compared to WT rats **(Supplementary figure 3H-G)**. In the spleen, TCR^-^ cells showed similar proportion of CD45RC^high^ and CD45RC^low/-^ cells in *Dmd*^*mdx*^ and WT animals **(Supplementary figure 3F-H)**.

WT and *Dmd*^*mdx*^ rats were injected with the same anti-CD45RC MAb used in the transplantation model described above ^14^ from week 2, since at this time point the leukocyte infiltration into the muscle has not yet appeared, and every 3.5 days and up to week 12 when grip force and mononuclear cells from muscle and spleen were analyzed.

At 12 weeks of age, treatment with anti-CD45RC MAb significantly depleted CD8^+^CD45RC^high^ T cells in both muscle and spleen of *Dmd*^*mdx*^ and in spleen of WT rats whereas CD8^+^CD45RC^low/-^ T cells were unchanged in both muscle and spleen **(Fig. 6A)**. Numbers of CD4^+^CD45RC^high^ T cells in the spleen were decreased but it did not reach statistical significance **(Fig. 6B)**. CD4^+^CD45RC^low/-^ **(Fig. 6B)** and FoxP3^+^ CD4+ T cells (data not shown) were maintained in both muscles and spleen. As in the transplantation models, other leukocytes that are CD45RC^high^ and TCR^-^, such as macrophages and B cells were not depleted by anti-CD45RC treatment **(Fig. 6C)**.

**Figure 6.**
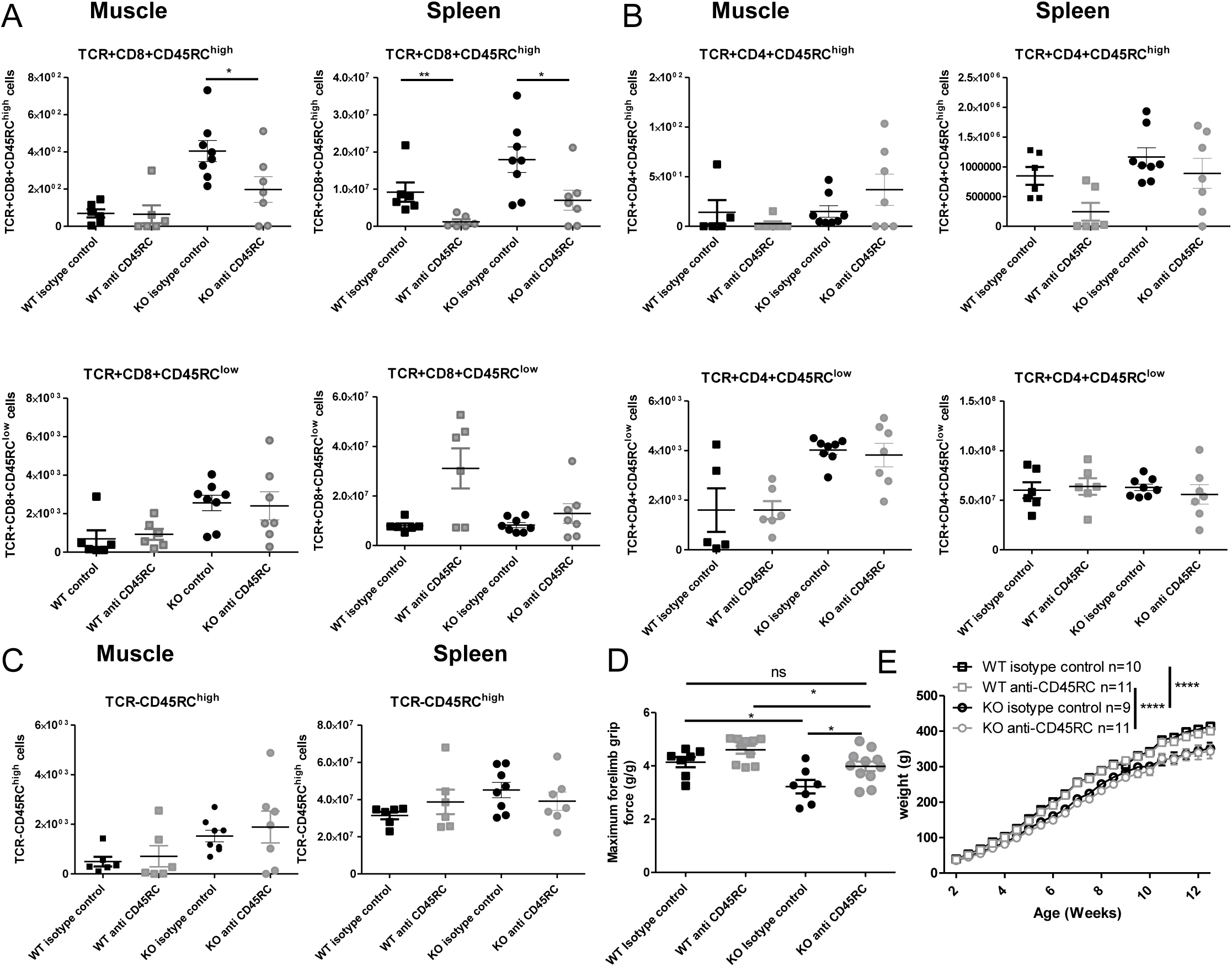
Effect of treatment with anti-CD45RC on lymphoid cell populations, forelimb muscle strength and animal growth. Hind limb muscles or spleen from WT or *Dmd*^*mdx*^ (KO) rats were harvested at 12 weeks of age, collagenase digested and analyzed by cytofluorimetry. **A)** Total numbers of viable CD45^+^TCR^+^CD8^+^CD45RC^high^ cells (upper panels) or viable CD45^+^TCR^+^CD8^+^CD45RC^low/-^ (lower panels) cells per gram of skeletal muscle (left panels) and of total spleen (right panels). **B)** Total numbers of viable CD45^+^TCR^+^CD4^+^CD45RC^high^ cells (upper panels) or viable CD45^+^TCR^+^CD4^+^CD45RC^low/-^ (lower panels) cells per gram of skeletal muscle (left panels) and of total spleen (right panels). **C)**. Total numbers of viable CD45^+^TCR^-^ CD45RC^high^ cells per gram of skeletal muscle (left panels) and of total spleen (right panels). **D)** Muscle strength in *Dmd*^*mdx*^ rats after treatment with an anti-CD45RC MAb. Wild-type (WT) or *Dmd*^*mdx*^ rats received intraperitoneal injections of the anti-rat CD45RC MAb (clone OX22, 1,5 mg/kg, every 3.5 days) or isotype control Mab (clone 3G8, 1,5 mg/kg, every 3.5 days) from week 2 to week 12 of age when muscle strength was analyzed using a grip test. Each point represents a single animal analyzed in two different experiments. * p< 0.05. Results were obtained from several experiments performed using all groups of animals in each experiment. **E)** Weight curves for animal growth were determined serially. ****p<0.001 between *Dmd*^*mdx*^ and WT rats for both treatments but no difference between *Dmd*^*mdx*^ rats treated with anti-CD45RC vs. isotype control.

At week 12 the animals were analyzed using a grip test. As previously reported ^3^, *Dmd*^*mdx*^ rats had a 30% reduction in forelimb strength compared to WT littermates **(Fig. 6D).** The treatment with anti-CD45RC MAb significantly improved muscle strength in *Dmd*^*mdx*^ treated rats vs. *Dmd*^*mdx*^ control animals **(Fig. 6D).** Furthermore, values for *Dmd*^*mdx*^ rats treated with anti-CD45RC MAb were indistinguishable of those of littermate WT controls **(Fig. 6D),** despite that they showed a significantly lower strength vs. WT animals treated with anti-CD45RC, but this was due to a slight non-significantly increase in muscle strength of WT animals treated with anti-CD45RC vs. WT isotype control-treated animals **(Fig. 6D).** The weight gain curves of *Dmd*^*mdx*^ animals were lower as compared to WT animals and treatment with anti-CD45RC neither modified this curve (**Fig. 6E**), nor the general aspect of the skeletal muscle fibrosis **(Supplementary figure 4A-B)** and CK levels in serum **(Supplementary figure 4C)**.

Thus, anti-CD45RC treatment resulted in increased muscle strength in *Dmd*^*mdx*^ rats and it was associated to depletion of T CD8^+^CD45RC^high^ cells.

### Treatment with prednisolone improved skeletal muscle strength but had secondary effects

Since CS are standard treatment in DMD patients ^6^, we analyzed the clinical effect of prednisolone on muscle strength of *Dmd*^*mdx*^ rats as well as in immune cells in skeletal muscle and spleen of *Dmd*^*mdx*^ rats.

Prednisolone-treated rats also showed at 12 weeks of age a significant decrease of CD8^+^CD45RC^high^ T cells in both muscle and spleen of *Dmd*^*mdx*^ rats and in spleen of WT rats **(Fig. 7A).** CD8^+^CD45RC^low/-^ T cells were maintained **(Fig. 7A).** CD4^+^CD45RC^high^ T cells were significantly decreased in spleen but not in muscle **(Fig. 7B).** CD4^+^CD45RC^low^ T cells were also decreased in the spleen but not in the muscle **(Fig. 7B).** Other leukocytes that are CD45RC^high^ and TCR^-^, such as macrophages and B cells were not depleted by prednisolone treatment **(Fig. 7C)**.

**Figure 7.**
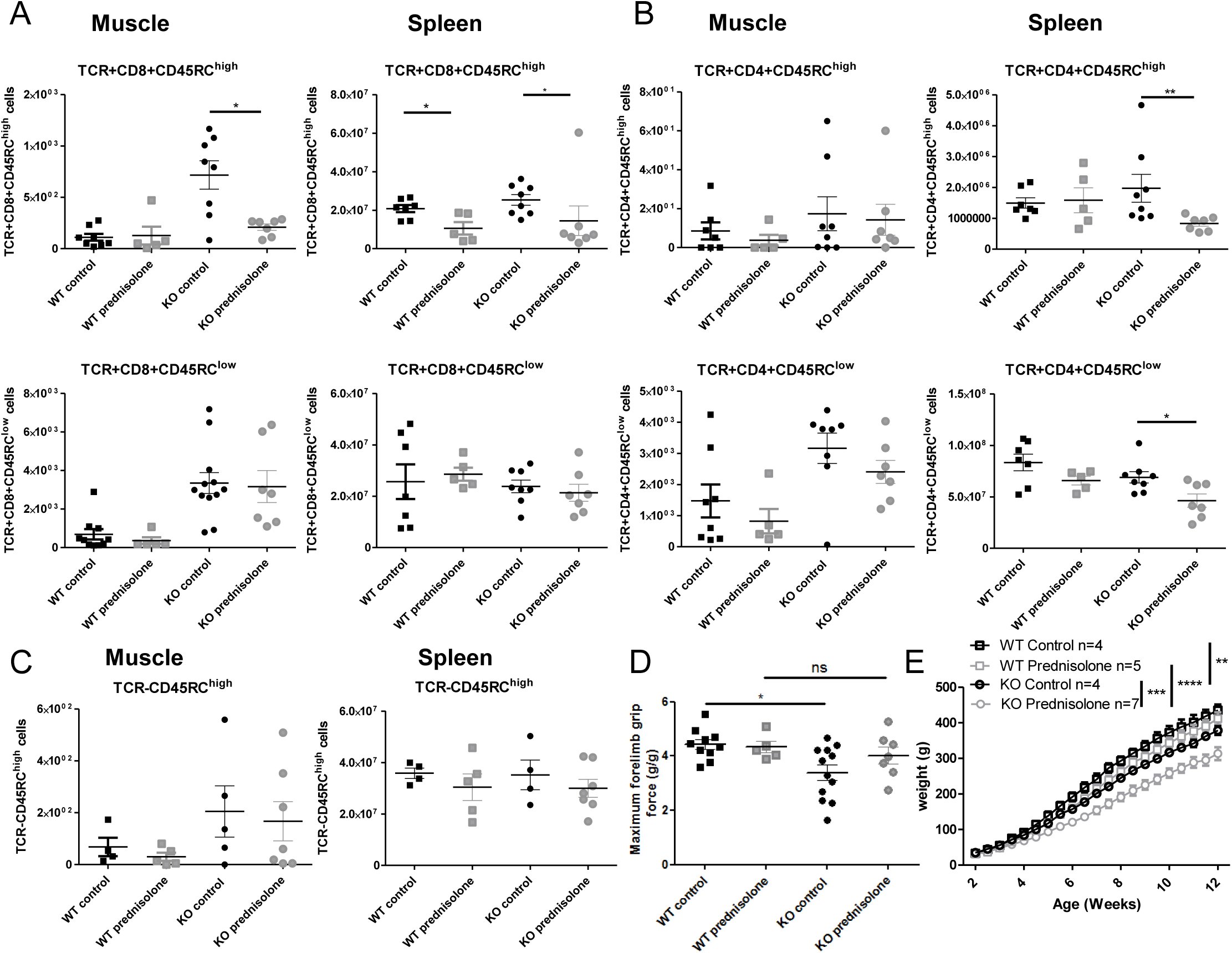
Treatment with prednisolone on lymphoid cell populations and forelimb muscle strength. Wild-type (WT) or *Dmd*^*mdx*^ (KO) rats received from week 2 of age intraperitoneal injections of prednisolone (0.5 mg/kg, 5 days per week) or NaCl up to week 12. **A)** Hind limb muscles or spleen from WT or *Dmd*^*mdx*^ were harvested, collagenase digested and analyzed by cytofluorimetry. Total numbers of viable CD45^+^TCR^+^CD8^+^CD45RC^high^ cells (upper panels) or viable CD45^+^TCR^+^CD8^+^CD45RC^low/-^ (lower panels) cells per gram of muscle (left panels) and of total spleen (right panels). * p< 0.05. **B)** Total numbers of viable CD45^+^TCR^+^CD4^+^CD45RC^high^ cells (upper panels) or viable CD45^+^TCR^+^CD4^+^CD45RC^low/-^ (lower panels) cells per gram of muscle (left panels) and of total spleen (right panels). **C)** Total numbers of viable CD45^+^TCR^-^ CD45RC^high^ cells per gram of skeletal muscle (left panels) and of total spleen (right panels). **D)** Muscle strength was analyzed using a grip test. Each point represents a single animal analyzed in two different experiments. * p< 0.05. Results were obtained from several experiments performed using all groups of animals in each experiment. **E)** Weight curves for animal growth were determined serially. **<0.01 and ****<0.0001 for *Dmd*^*mdx*^ and WT with NaCl and prednisolone but importantly ***p<0.001 between *Dmd*^*mdx*^ rats NaCl vs. prednisolone.

Simultaneously, *Dmd*^*mdx*^ rats treated with prednisolone showed significantly increased muscle strength at 12 weeks to levels identical to those of WT or anti-CD45RC-treated rats **(Fig. 7D).** Prednisolone-treated *Dmd*^*mdx*^ rats showed marked secondary effects, as shown by a severe (25%) and significant reduction in growth as compared to WT rats but also to NaCl-treated *Dmd*^*mdx*^ rats (**Fig. 7E**). Prednisolone had no effect on the growth of WT animals (**Fig. 7E**). Muscle tissue fibrosis **(Supplementary figure 4A-B)** and CK levels in serum **(Supplementary figure 4C)** were not modified by prednisolone treatment.

Thus, as compared to anti-CD45RC treatment, prednisolone also increased muscle strength but showed a larger decrease in cell populations including not only CD8^+^CD45RC^high^ cells in muscle and spleen, but also CD4^+^CD45RC^high^ and CD4^+^CD45RC^low^ cells in spleen and had a strong negative effect on the growth of *Dmd*^*mdx*^ animals.

### Presence of T CD45RC^high^ cells in skeletal muscles and blood of DMD patients

To further explore the potential of CD45RC as an immunotherapeutic target, we evaluated the presence of CD45RC^high^ cells in peripheral blood in DMD patients. Cytofluorimetry analysis showed the presence of CD45RC^high^ and CD45RC^low/-^ among both CD4^+^ or CD8^+^ T cell compartments in blood of DMD patients in proportions comparable to those of age-matched young individuals hospitalized for pathologies not involving the immune system or other neuromuscular diseases **(Supplementary figure 5).** As for young controls, B and NK cells from DMD patients were all CD45RC^high^ whereas monocytes and PMN were all CD45RC^-^ **(Supplementary figure 6)**.

Furthermore, the presence of CD45RC brightly positive cells was confirmed in muscle biopsies from DMD patients and not of normal individuals, as it was the case in muscles of *Dmd*^*mdx*^ vs. WT animals **(Fig. 8).**

**Figure 8.**
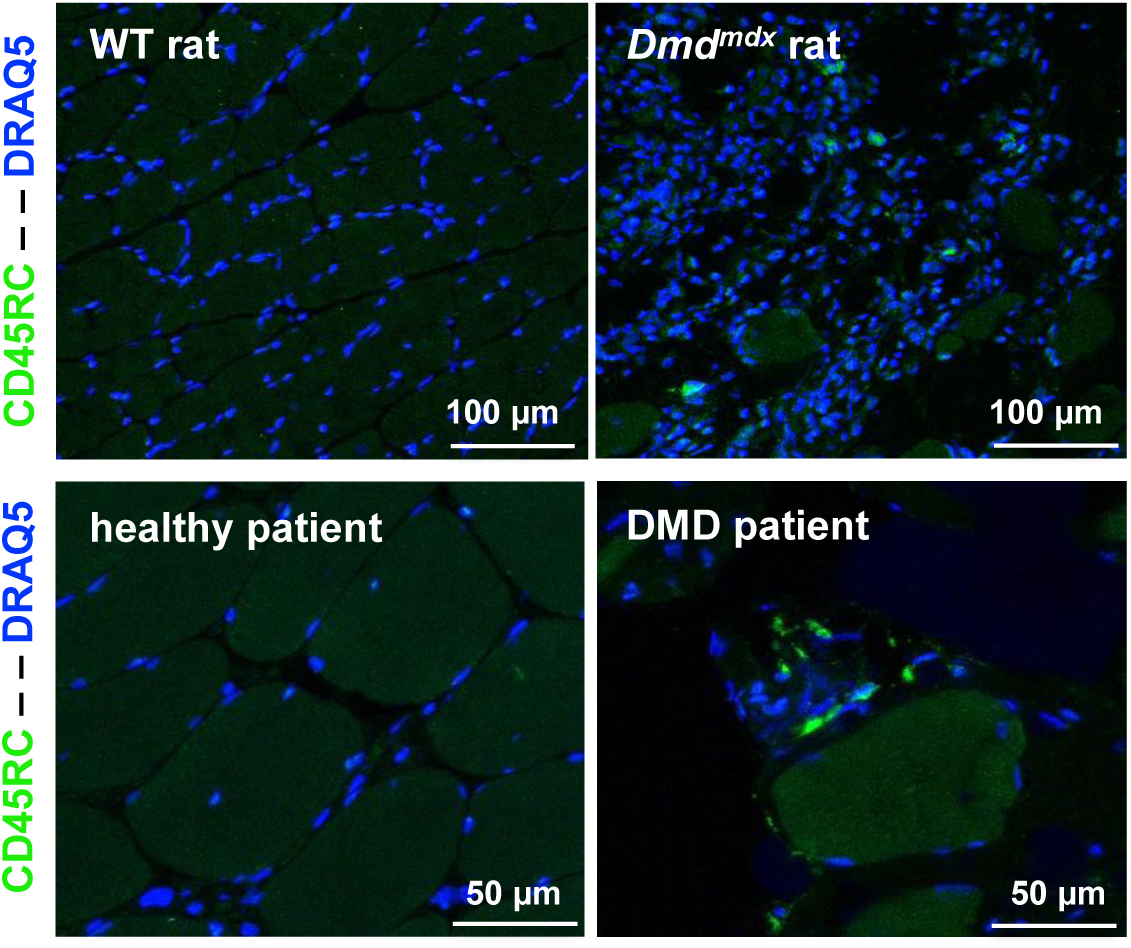
CD45RC^+^ cells in rat and human dystrophin-deficient skeletal muscles. Skeletal muscle samples from rat (*Biceps femoris*) and humans (*Paravertebralis*), either from dystrophin deficient individuals (n=2) or without muscle pathology (n=2). Pictures are representative images of frozen tissue sections probed with Draq5 to label nuclei and with anti-rat or human anti-CD45RC MAbs (green).

## Discussion

DMD patients and *mdx* mice show muscle infiltration by different types of leukocytes and production of a variety of mediators that have been shown to play facilitating or protecting roles in the evolution of the disease ^5^. Not only inflammation and innate immune responses are present, but also adaptive immune responses including anti-dystrophin T cells and Treg cells are involved in DMD patients ^8, 10^ and mdx mice ^5, 11, 12^. CS are one of the only standard treatments that DMD patients receive and that prolong ambulation by about 2 years. Nevertheless, increase muscular strength responses are variable, incomplete and always associated to serious side effects ^6, 7^. Despite that the precise mechanisms of action of CS in DMD patients are ill defined, anti-inflammatory effects are likely important ^6, 7^. Thus, there are unmet clinical needs to treat the inflammatory and immune effects caused by dystrophin deficiency while awaiting for curative gene or stem cell therapies. It is even very likely that these immunotherapies will be associated to gene and cell therapy to inhibit immune responses against the vectors, transgene products or antigenic cellular products.

The *mdx* mouse is a very useful model but fails to reproduce key symptoms of DMD patients such as muscular weakness ^2^. Thus, although several immunotherapies were successful in *mdx* mice, such as intravenous immunoglobulin ^18^, anti-TNFα antibodies ^19^, IL-6 blocking ^20^, tranilast ^21^, heme oxygenase-1 (HO-1) inducers ^22^, IL-1 receptor antagonist ^23^ and IL-2 complexes to amplify CD4^+^ Tregs ^12^, their potential effect in DMD patients is uncertain. *Dmd*^*mdx*^ rats reproduce skeletal and cardiac muscular weakness at early time points and develop skeletal and cardiac muscle tissue lesions that resemble those observed in DMD patients ^3, 4^. In the present manuscript we describe that *Dmd*^*mdx*^ rats present mononuclear cells infiltrating both skeletal and cardiac muscles that appeared early, between 2 and 4 weeks of age, and that had greatly decreased by 16 weeks of age. The majority of these mononuclear cells were CD68^+^ and SIRPα^+^ macrophages and the proportion of M2 CD163^+^ increased with time. Macrophages appear early in both *mdx* mice (2 weeks) and DMD patients (2-year-old) ^24^. M2 macrophages have been shown to play protective and regenerative roles in early stage disease in *mdx* mice ^5^. CD4^+^ and CD8^+^ T cells, including Tregs, were also increased in muscles of *Dmd*^*mdx*^ rats compared to controls. The lesions of muscular fibers, as analyzed by CK levels in serum, followed the kinetics of leukocyte infiltration, with normal levels at 2 weeks of age and a peak between 4 and 8 weeks of age for a later decrease, possibly reflecting a more pronounced immune attack at early rather than late time points.

Cytokines were produced at increased levels by mononuclear cells from *Dmd*^*mdx*^ rats compared to controls at 8 and/or 12 weeks of age, including IL-1β and TNFα. These cytokines are increased in DMD patients and mdx mice, have been described as potential immunotherapy targets ^25^, since anti-TNFα treatment reduces early muscle damage in *mdx* mice ^19^ could be targeted in the future in *Dmd*^*mdx*^ rats. Several anti-inflammatory molecules, such as HO-1, IL-10 and TGFβ, as well as the muscle trophic factor amphiregulin ^11^, were also produced, likely as a response to inflammation and ongoing immune responses, as previously described in mdx mice and DMD patients ^5^. Inhibition of TGFβ has been shown to play a dual role since early neutralization in mdx mice decreases fibrosis but increases T cell infiltration and inflammation^26^.

We have recently shown that treatment with an anti-CD45RC MAb in a rat model of heart allograft rejection could induce permanent allograft acceptance ^14^. Furthermore, anti-CD45RC MAb treatment prevented GVHD in immune humanized NSG (NOD *Scid* Gamma) mice ^14^. Anti-CD45RC treatment depleted T cells that were CD45RC^high^, comprising naïve T cells, precursors of Th1 cells and T effector memory cells including TEMRA cells, whereas CD8^+^ and CD4^+^ Tregs both in rats and humans are CD45RC^low/-13, 27^ and thus were spared. These CD45RC^low/-^ CD8^+^ and CD4^+^ Tregs that were specific of donor alloantigens could impose allograft tolerance in newly grafted irradiated recipients following adoptive cell transfer. Importantly, immune responses against third party donors and exogenous antigens were preserved during treatment with anti-CD45RC, thus depletion of CD45RC^high^ cells does not inhibit all immune responses.

We thus reasoned that treatment of *Dmd*^*mdx*^ rats with anti-CD45RC MAbs could eliminate CD45RC^high^ T effector cells and their precursors and enrich CD45RC^low/-^ Tregs that could then act at the same time, not only as inhibitors of immune responses by CD45RC^high^, but also favor tissue repair and homeostasis by CD45RC^low/-^, such as it has been described for CD4^+^ Tregs both in muscle ^11^ and adipose tissue ^28^. In the present manuscript we show that anti-CD45RC treatment improved muscle strength to the levels of WT animals and that this was associated to a depletion of CD8^+^ CD45RC^high^ T cells at 12 weeks. CD4^+^ CD45RC^high^ T effector cells were decreased but not significantly at this time of treatment. Although CD45RC^low/-^ CD8^+^ or CD4^+^ Tregs were not numerically increased in *Dmd*^*mdx*^ rats this was also the case in rats that were tolerant to transplanted organs after anti-CD45RC MAb treatment ^14^. Whether their function is increased and play a role in the amelioration of muscle strength observed in these animals remains to be analyzed in future studies.

As for the anti-CD45RC treatment, corticosteroids resulted in a similar increase in muscular strength that was associated surprisingly to a specific decrease in CD8^+^ CD45RC^high^ T cells in muscle but also to a more widespread decrease of CD4^+^CD45RC^high^ and CD4^+^CD45RC^low/-^ cells in spleen. This effect has also been observed in DMD patients treated with steroids ^5^. DMD patients treated with corticosteroids showed decreased T cells against dystrophin ^10^.

The secondary effects of steroids were observed in *Dmd*^*mdx*^ rats, whereas anti-CD45RC treated animals did not show obvious clinical abnormalities and no weight loss. Since patients suffer from several important side effects of steroids, anti-CD45RC treatment could result in similar muscle improvement than corticosteroids but without side effects. A potential side effect of anti-CD45RC treatment could be generalized immunosuppression but we have already demonstrated that rats treated with anti-CD45RC treatment could mount normal primary immune responses to new antigens as well as memory immune responses after secondary immunization ^14^.

MAbs against CD45RA ^25^, CD45RO/B ^29^ and CD45RB ^30^ have been used to treat organ rejection and/or GVHD but none has been used as isolated treatment, neither in animal models of DMD, nor on animal models of muscle lesions. Even if anti-CD45RA or antiCD45RO/B could be used in the future, since between 50 to 90% of both of the CD8^+^ and CD4^+^ Tregs are CD45RA^high^ and CD45RB^high 14^, the outcome of treatment with anti-CD45RC is clearly targeting different cell populations and thus distinct and likely more favorable since it preserves Tregs. Although in mdx mice depletion of total CD4^+^ or CD8^+^ cells ameliorates histopathology ^31^, none of other tolerizing treatments based on MAbs and used in organ transplantation, GVHD or autoimmunity, such as anti-CD3, anti-CD127, anti-CD28, have been previously used in models of DMD and thus the results in the present manuscript could stimulate the use of these other reagents.

## Materials and methods

### Animal experiments and ethical aspects

*Dmd*^*mdx*^ rats have been previously described ^3^. *Dmd*^*mdx*^ and wild-type littermate were raised in SPF conditions. All the animal care and procedures performed in this study were approved by the Animal Experimentation Ethics Committee of the Pays de la Loire region, France, in accordance with the guidelines from the French National Research Council for the Care and Use of Laboratory Animals (Permit Numbers: CEEA-PdL-10792 and CEEA-PdL-8986). All efforts were made to minimize suffering. The rats were housed in a controlled environment (temperature 21±1°C, 12-h light/dark cycle). Blood samples from 2 DMD patients were obtained as part of their standard care management in the hospital and after obtaining informed consent from both patients and their parents. Control blood samples were collected from children who had been admitted in Nantes University Hospital without immune deficiency. The biocollection used for this analysis is the “pediatrics” collection (Ref: MESR DC-2011-1399) which is a prospective monocentric collection managed by the University Hospital of Nantes and approved by the local ethics committee. None of the legal representatives of the children objected to let them take part in this biocollection. Tissue samples were obtained from the *Paravertebralis* muscle of four 12 year-old patients (two DMD patients and two patients free of known muscular disease). Patients were operated at the Department of Pediatric Surgery of the Centre Hospitalier Universitaire (CHU) de Nantes (France). They gave written informed consent. All protocols were approved by the Clinical Research Department of the CHU (Nantes, France), according to the rules of the French Regulatory Health Authorities (Permit numbers: MESR/DC-2010-1199). Biological sample bank was constituted in compliance with the national guidelines regarding the use of human tissue for research (Permit numbers: CPP/29/10).

### Preparation of muscle and spleen single-cell suspensions

Muscles of both hindlimbs from WT or *Dmd*^*mdx*^ rats were excised without adipose tissue, rinsed with PBS and weighed. Muscles were minced, placed in gentleMACS C tubes (Miltenyi Biotec) with collagenase D (4ml/g of muscle), dissociated using the gentleMACS^TM^ dissociator (gentleMACS program “m_muscle_01”) and incubated for two runs of 30 min at 37°C under continuous rotation. After the initial run, undigested muscle was filtrated on a mesh strainer and the resulting cell suspension was centrifuged and resuspended in PBS FCS 2% 1 mM EDTA. The remaining undigested muscle was further incubated in fresh collagenase for a new run of 30 min. The debris-free cell suspensions were centrifuged, resuspended in PBS FCS 2% 1 mM EDTA, and pooled with cells from the first digestion. Pooled cells were then applied to 15 ml Histopaque 1077density gradient (Eurobio) and centrifuged at 1000 x g for 30min. The cells at the interface were collected, washed, resuspended in PBS FCS 2% 1mM EDTA and counted.

Spleen was harvested, perfused with collagenase D, minced and incubated for 15 min at 37°C. Spleen fragments were then scraped in the presence of PBS FCS 2% 1mM EDTA and mononuclear cells were recovered using a density gradient (Histopaque 1077, Eurobio). The cells at the interface were collected, washed, resuspended in PBS FCS 2% 1mM EDTA and counted.

### Staining of rat cells for flow cytometry analysis

Cytofluorimetry analysis was performed as previously described in detail ^32^. Briefly, single-cell suspensions from muscle or spleen were stained with MAbs against the following antigens: CD45 as a pan leukocyte (clone OX-1), TCRαβ (clone R7/3), CD45RA in B cells (clone OX33), CD45R/B220 in B cells (clone His24), anti-granulocytes (RP-1 and His48), CD4 (clone w3/25), CD45RC (clone OX22 or clone OX32), CD25 (clone OX39), CD8 (clone OX8), CD172a/SIRPα (clone OX41), CD161 in NK and myeloid cells (clone 3.2.3), CD163 in macrophages (clone ED2), CD68 for macrophages (clone ED1) and with viability dye eFluor506 or eFluor450 from eBiosciences to assess cell viability. Analysis was performed on a BD FACS Verse with FACSuite Software version 1.0.6. Post-acquisition analysis was performed with FlowJo software.

### Serum creatinine phosphokinase and cytokine levels

Blood was collected under anesthesia, serum was isolated and immediately frozen at −20°C. Total creatinine phosphokinase (CK) activity was determined in the biochemistry department of Nantes University Hospital.

Levels of IL-1β, IL-6, IL-10 and TNFα in the serum of *Dmd*^*mdx*^ or WT littermate rats, were measured by multiplex assay (Luminex technology) (R&D systems) following to the manufacturer’s instructions.

### Quantitative RT-PCR

Quantification of mRNA levels was performed as previously described in detail ^33^. Briefly, total RNA extraction has been performed on mononuclear cells from skeletal muscles using RNeasy Mini Kit (Qiagen) according to the manufacturer’s instructions. Then quantification and quality analysis were done on Caliper LabChip GX II (PerkinElmer). RNA with a quality score between 7 and 10 were retro-transcribed using oligo-dT and M-MLV reverse transcriptase (Life Technologies). Fast SybrGreen Master Mix 2x was used to performed qPCR on ViiA 7 (Applied Biosystems) on cDNA in duplicate for each target according to the manufacturer’s instructions. qPCR reaction conditions were 20 seconds at 95°C followed by 40 cycles of 1 second at 95°C, 20 seconds at 60°C and 20 seconds at target melting temperature minus 3°C, ended by a melting curve stage. Calculations were made by DDCt method. The primers used in this study are listed in **table 1**.

**Table 1.**
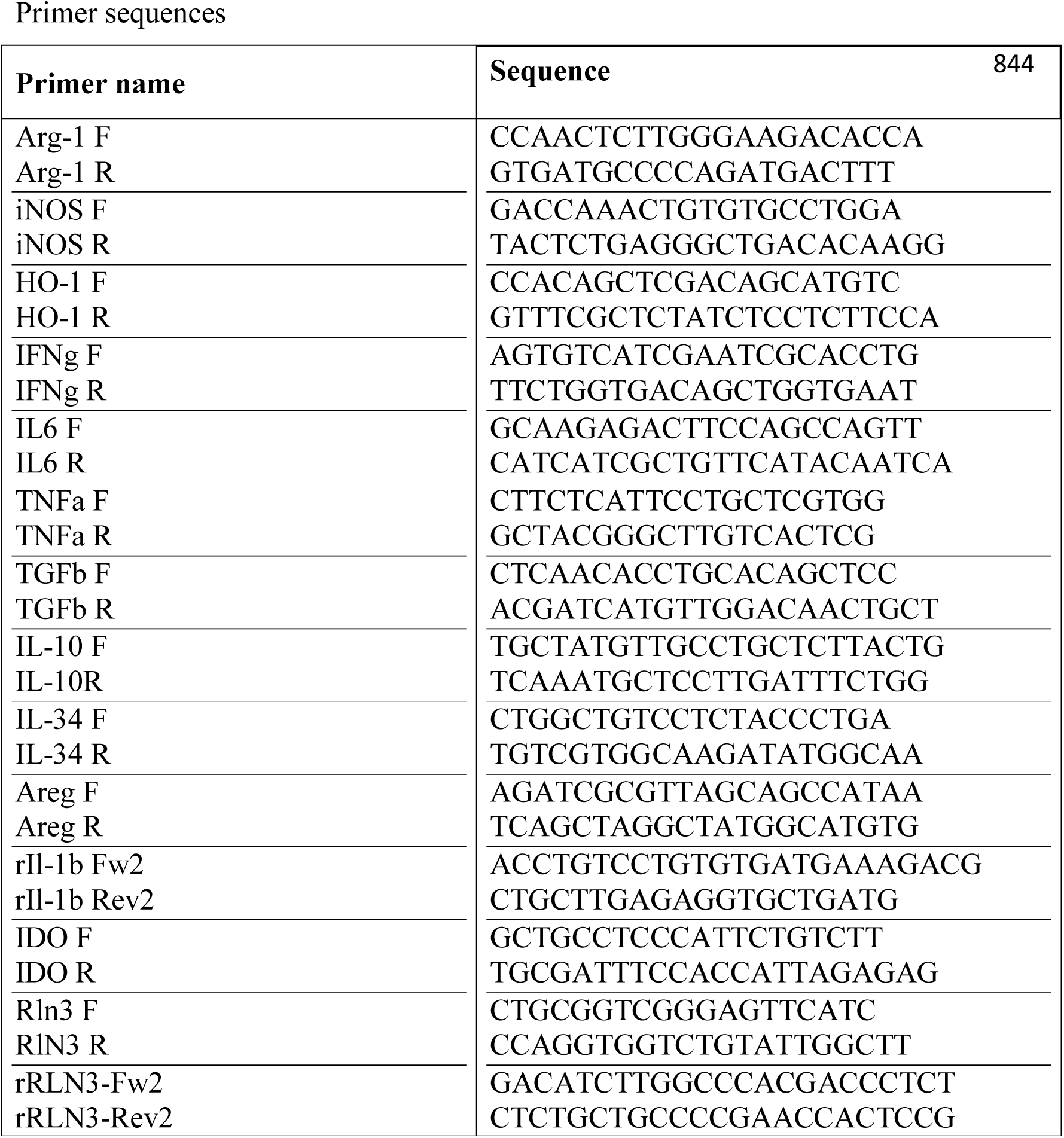
Primer sequences

### Immunohistological analysis and fibrosis quantification

Immunohistochemistry was performed as previously described in detail ^3^. Briefly, tissue samples of *Biceps femoris* and cardiac ventricular muscles were harvested at 8 and 12 weeks of age and frozen and 8-μm-thick sectioned for immunofluorescence labelling. Sections were preliminary fixed with acetone for CD3 labelling and with acetone (30%) in methanol for CD68, CD163 and CD45RC labelling (10 min, room temperature) and incubated with 0.2% triton in PBS (10 min, room temperature). Sections were then blocked with 10% goat serum in PBS and incubated with the primary antibodies. Rabbit polyclonal antibody for CD3 (DakoCytomation, Glostrup, Denmark), mouse monoclonal antibodies for rat CD68, CD163 and CD45RC were used respectively at 1:50; 1:200 and 1:200 (overnight, 4°C). After washing, goat anti-rabbit and goat anti-mouse antibodies coupled with Alexa 488 (InVitroGen, Carlsbad, CA) were respectively used to reveal CD3 and CD68 primary antibody (1 h, room temperature). Section were incubated with wheat germ agglutinin Alexa Fluor 555 conjugate for connective tissue labelling (Molecular Probes, Eugene, OR) diluted 1:700 in PBS (overnight, 4°C) and nuclei were then labelled with Draq5 (BioStatus Ltd, Shepshed, UK) diluted at 1:1000 (10min, room temperature). Immunofluorescence labeling was analyzed with a laser scanning confocal microscope (Zeiss, LSM880, Jena, Germany).

For human muscle, biopsies were obtained from DMD patients undergoing surgery for spinal deformities and from young individuals undergoing muscle biopsy for other diagnosis. Tissue was frozen, sectioned and processed as described above for rat tissue using an anti-human CD45RC MAb (BD Biosciences).

### Treatment with anti-CD45RC and prednisolone

WT and *Dmd*^*mdx*^ rats received intraperitoneal injections of the anti-rat CD45RC MAb (clone OX22, mouse IgG1) or an isotype control MAb (clone 3G8, mouse IgG1) at 1.5 mg/kg, every 3.5 days from week 2 to week 12 of age as previously described ^14^. Prednisolone was administered by daily intraperitoneal injections of 0.5 mg/kg, close to the dose of 1 mg/kg in mdx mice ^34^ and 0.75 mg/kg in DMD patients ^35^ from week 2 to week 12 of age.

### Grip test

Grip test was performed as previously described in detail ^3^. Rats were placed with their forepaws on a grid and were gently pulled backward until they released their grip, as previously described. A grip meter (Bio-GT3, BIOSEB, France), attached to a force transducer measured the peak force generated.

### Statistical analysis

Mann–Whitney t test was used to compare numbers of cells in muscle and spleen of WT vs *Dmd*^*mdx*^, CK and cytokine levels in sera.

Two-way ANOVA test was used to compare growth curves.

---

**Supplementary figure 1. Number of SIRP**α**^+^ macrophages in skeletal muscle and spleen of *Dmd*^*mdx*^ rats.** Hind limb muscles and spleen were harvested from littermate wild-type (WT) or *Dmd*^*mdx*^ (KO) rats at the indicated time points of age. Muscle and spleen were digested with collagenase, mononuclear cells were isolated using a density gradient and analyzed by cytofluorimetry. **A)** Number of viable CD45^+^TCR^-^ CD45RA^-^SIRPα^+^ cells per gram of muscle (left panel) or whole spleen (right panel) at different time points. WT, n=4, 5, 7, 7, 9 at 2, 4, 8, 12 and 16 weeks, respectively; *Dmd*^*mdx*^, n=3, 6, 10, 11, 16 at 2, 4, 8, 12 and 16 weeks, respectively. ** p< 0.01, and *** p< 0.001. **B)** Representative dot-plot analysis of viable SSC CD45^+^TCRCD45RA^-^SIRPα^+^ cells mononuclear leukocytes from muscle (left panels) or spleen (right panels) from animals at 12 weeks of age.

**Supplementary figure 2. CK in sera of *Dmd*^*mdx*^ rats.** CK levels were determined simultaneously in all samples. WT (n=8, 6, 6, 9, 5 at 2, 4, 8, 12 and 16 weeks, respectively), *Dmd*^*mdx*^ (n=5, 4, 8, 8, 6 at 2, 4, 8, 12 and 16 weeks, respectively). * p<0.05.

**Supplementary figure 3. Expression profiles of CD45RC in different mononuclear cell populations**. Hind limb muscles and spleen were harvested from littermate wild-type (WT) or *Dmd*^*mdx*^ (KO) rats at the indicated time points of age. Muscle and spleen were digested with collagenase, mononuclear cells were isolated using a density gradient and analyzed by cytofluorimetry. **A)** Absolute numbers of CD45^+^CD45RA^-^TCR^+^CD8^+^CD45RC^high^ cells in muscle or spleen of WT or *Dmd*^*mdx*^ rats during time. **B)** Representative dot-plot analysis of viable CD45^+^CD45RA^-^TCR^+^CD8^+^CD45RC^high or low/-^ mononuclear leukocytes from muscle (left panels) or spleen (right panels) of WT or *Dmd*^*mdx*^ rats from animals at 12 weeks of age. **C)** Absolute numbers of CD45^+^CD45RA^-^TCR^+^CD4^+^CD45RC^low/-^ cells in muscle of spleen of WT or *Dmd*^*mdx*^ rats during time. **D)** Representative dot-plot analysis of viable CD45^+^CD45RA^-^TCR^+^CD4^+^CD45RC^high or low/-^ mononuclear leukocytes from muscle (left panels) or spleen (right panels) of WT or *Dmd*^*mdx*^ rats from animals at 12 weeks of age. **E)** Absolute numbers of CD45^+^CD45RA^-^TCR^+^CD4^+^CD45RC^high^ cells in muscle of spleen of WT or *Dmd*^*md*^ rats during time. **F)** Absolute numbers of CD45^+^TCR^-^CD45RC^low/-^ cells in muscle of spleen of WT or *Dmd*^*md*^ rats during time. **G)** Representative dot-plot analysis of viable CD45^+^TCR^-^CD45RC^high or low/-^ mononuclear leukocytes from muscle (left panels) or spleen (right panels) from animals at 12 weeks of age. **H)** Absolute numbers of CD45^+^TCR^-^ CD45RC^high^ cells in muscle of spleen of WT or *Dmd*^*mdx*^ rats during time.

**Supplementary figure 4. Effects of anti-CD45RC or prednisolone treatments on muscle fibrosis and serum CK levels.** Littermate wild-type (WT) or *Dmd*^*mdx*^ (KO) rats were treated with anti-CD45RC or prednisolone since week 2 of age. **A)** *Biceps femoris* muscles were harvested at 12 weeks of age, fixed and paraffin embedded, connective/fibrotic tissue was stained with picrosirius for connective tissue and the stained surface was quantified and expressed as the percentage of total area of the tissue analyzed (47 mm^2^). WT isotype control, n=6; WT anti-CD45RC, n=3; KO isotype control, n=6; KO anti-CD45RC, n=3; KO prednisolone, n=6. * p<0.05 vs. WT controls and anti-CD45RC-treated animals. **B)** Representative picrosirius (purple) staining for animals of the indicated group treatments. **C)** (left panel) Sera of *Dmd*^*mdx*^ and WT rats treated with prednisolone or vehicle (NaCl) were collected at 12 weeks of age and CK levels were determined simultaneously in all samples. WT NaCl, n=3; WT prednisolone, n=3; KO NaCl, n=5; KO prednisolone, n=6. (right panel) Sera of *Dmd*^*mdx*^ and WT rats treated with anti CD45RC or isotype control were collected at 4, 8 12 and 16 weeks of age and CK levels were determined simultaneously in all samples. WT isotype control (n=12, 4, 11, 4 at 4, 8, 12 and 16 weeks of age, respectively); WT anti CD45RC (n=13, 8, 11, 3 at 4, 8, 12 and 16 weeks of age, respectively); KO isotype control (n= 6, 4, 14, 3 at 4, 8, 12 and 16 weeks of age, respectively); KO anti CD45RC (n=8, 5, 13, 4 at 4, 8, 12 and 16 weeks of age, respectively).

**Supplementary figure 5. Cytofluorimetry analyses of CD45RC expression in blood T cells of DMD patients and controls.** Human peripheral blood was drawn, red blood cells were lysed and white blood cells were incubated with a viability dye and MAbs directly coupled with the indicated fluorochromes defining CD3, CD4, CD8 and CD45RC or isotype controls followed by cytofluorimetry analyzes. Ordinate depict reactivity with anti-CD45RC MAb or isotype control and the boxes define CD45RC^high^ or CD45RC^low/-^ cells. Abscissa depict reactivity with anti-CD4 or CD8 MAbs among CD3^+^ cells. Controls were young patients (6-17 years-old) comparable in age to DMD patients and that were hospitalized for pathologies not involving the immune or the neuromuscular systems.

**Supplementary figure 6. Cytofluorimetry analyses of CD45RC expression in blood non-T cells of DMD patients and controls.** Human peripheral blood was drawn, red blood cells were lysed and white blood cells were incubated with a viability dye and MAbs directly coupled with the indicated fluorochromes defining CD14^+^ monocytes, CD19^+^ B cells, CD16+56^+^ NK cells and CD45RC^high^ cells or isotype controls followed by cytofluorimetry analyzes. Only one DMD and one control patients are showed as representative example. Ordinate depict reactivity with anti-CD45RC MAb or isotype control and the boxes define CD45RC^high^ cells. Abscissa depict reactivity with anti-CD14, anti-CD19 or anti-CD16+56 MAbs. Controls were young patients (6-17 years-old) comparable in age to DMD patients and that were hospitalized for pathologies not involving the immune or the neuromuscular systems.

Acknowledgments
This work was financially supported by the Région Pays de la Loire through Biogenouest, and the “TEFOR” project funded by the “Investissements d’Avenir” French Government program, managed by the French National Research Agency (ANR) (ANRII-INSB-0014). This work was realized in the context of the Labex IGO project (n°ANR-11-LABX-0016-01) and the IHU-Cesti project (ANR-10-IBHU-005) which both are part of the “Investissements d’Avenir” French Government program managed by the ANR. The IHU-Cesti project is also supported by Nantes Métropole and Région Pays de la Loire. This work was also realized in the context of the support provided by the Fondation Progreffe.

